# Same same but different: a web-based deep learning application for the histopathologic distinction of cortical malformations

**DOI:** 10.1101/804682

**Authors:** J. Kubach, A. Muehlebner-Farngruber, F. Soylemezoglu, H. Miyata, P. Niehusmann, M. Honavar, F. Rogerio, S-H. Kim, E. Aronica, R. Garbelli, S. Vilz, A. Popp, S. Walcher, C. Neuner, M. Scholz, S. Kuerten, V. Schropp, S. Roeder, P. Eichhorn, M. Eckstein, A. Brehmer, K. Kobow, R. Coras, I. Bluemcke, S. Jabari

## Abstract

We trained a convolutional neural network (CNN) to classify H.E. stained microscopic images of focal cortical dysplasia type IIb (FCD IIb) and cortical tuber of tuberous sclerosis complex (TSC). Both entities are distinct subtypes of human malformations of cortical development that share histopathological features consisting of neuronal dyslamination with dysmorphic neurons and balloon cells. The microscopic review of routine stainings of such surgical specimens remains challenging. A digital processing pipeline was developed for a series of 56 FCD IIb and TSC cases to obtain 4000 regions of interest and 200.000 sub-samples with different zoom and rotation angles to train a CNN. Our best performing network achieved 91% accuracy and 0.88 AUCROC (area under the receiver operating characteristic curve) on a hold-out test-set. Guided gradient-weighted class activation maps visualized morphological features used by the CNN to distinguish both entities. We then developed a web application, which combined the visualization of whole slide images (WSI) with the possibility for classification between FCD IIb and TSC on demand by our pretrained and build-in CNN classifier. This approach might help to introduce deep learning applications for the histopathologic diagnosis of rare and difficult-to-classify brain lesions.

## Introduction

Deep learning showed remarkable success in medical and non-medical image-classification tasks in the past 5 years [1–3], finding its way into applications for digital-pathology such as classification, cell detection and segmentation. Based on these tasks more abstract functions like disease grading, prognosis prediction and imaging biomarkers for genetic subtype identification have been established [4, 5]. Successful examples range from utilization in different types of cancer detection/classification/grading [6, 7], classification of liver cirrhosis [8], heart failure detection [9] and classification of Alzheimer plaques [10].

The most commonly used deep learning architectures are convolutional neural networks (CNN) (Figure 1c). CNN’s are assembled as a sequence of levels consisting of convolutional layers and pooling layers, followed by fully connected layers with a problem specific activation function in the end [11, 12]. Each convolutional layer consists of feature maps connected with an area of the previous layer and a set of specific weights for each feature map. The convolutional layer is followed by pooling layers, which compute the maximum or average of a group of feature maps. This pooling-operation merges related values into features and reduces the dimension by taking input from multiple overlapping feature maps. This combination enables the CNN to correlate groups of local values to detect patterns as well as making motifs invariant to the exact location in the images [11]. In other words, CNN will learn features in images without explicitly showing, segmenting or marking important features in a given motif.

**Figure 1.**
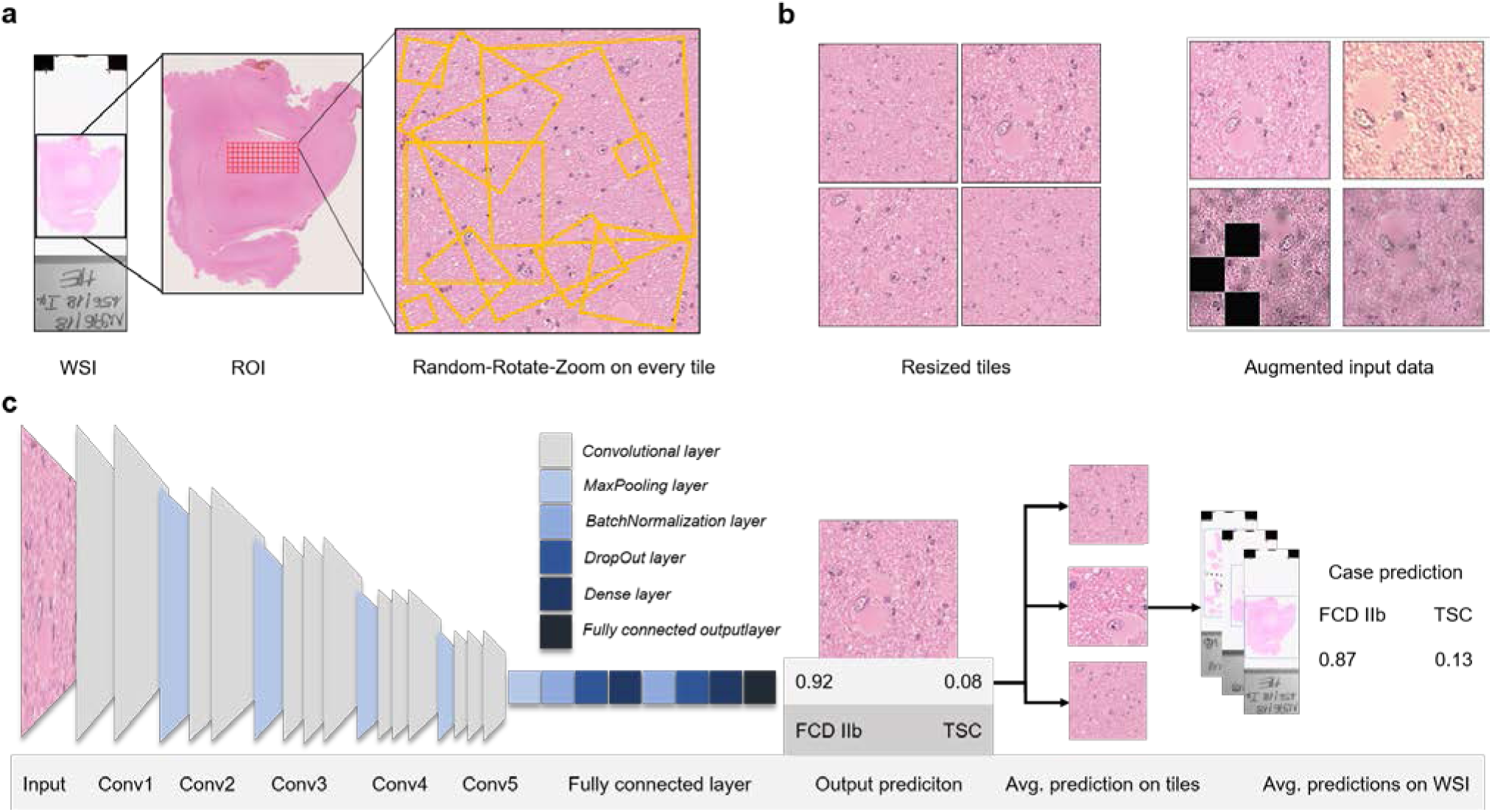
Workflow overview. **a|** Workflow beginning with digitized WSI then extracting the tiles of a ROI and 10 random-rotate-zoom example subtiles visualized on a single 2041×2041 pixel tile. **b|** Example tiles obtained through random-rotate-zoom and resizing then further augmenting one tile to show a range of augmentation techniques used. **c|** The unchanged VGG16 architecture merges into a custom top layer consisting of BatchNormalization, DropOut and Dense Layers merging into a fully connected output layer. The predictions on single tiles are averaged to obtain the prediction of a WSI and the WSI-predictions will be averaged to obtain the prediction of a case.

Malformations of Cortical Development (MCD) represent common brain lesions in patients with drug-resistant focal epilepsy and surgical resection is a beneficial treatment option [13, 14]. Amongst the many MCD conditions described in the literature, focal cortical dysplasia (FCD) and cortical tuber of tuberous sclerosis complex (TSC) share histopathological communalities difficult to distinguish at the microscopic level. TSC is a variable neurocutaneous disorder involving benign tumors and hamartomatous lesions in different organ systems, most commonly in the brain, skin and kidneys. TSC is caused by autosomal dominant mutations in the TSC 1 (hamartin) and TSC 2 (tuberin) gene. As a complex, TSC 1 and TSC 2 inhibit the mammalian target of rapamycin complex (mTORC1) [15, 16]. Also, TSC can be diagnosed clinically if two major features or one major and two minor features are present, following the international TSC diagnostic criteria [17]. FCDs are a heterogenous subgroup of MCD, which can be located throughout the cortex. FCDs can be classified using the three-tiered ILAE classification system, subdividing the FCDs based on histopathological findings including abnormal radial and tangential cortical lamination, dysmorphic neurons, balloon cells and adjacent to other principal lesions [18]. The subtype FCD IIb is histomorphologically characterized by dysmorphic neurons and balloon cells, disrupted cortical lamination as well as blurred boundaries between gray and white matter [18].

In routine histopathology work-up, TSC can hardly be distinguished from FCD IIb, in particular, balloon cells are not discernible from giant cells in TSC patients. Cortical tubers are also defined by disrupted cortical lamination, thus making a definite diagnosis based on the histomorphological findings difficult to obtain (Figure 2).

**Figure 2.**
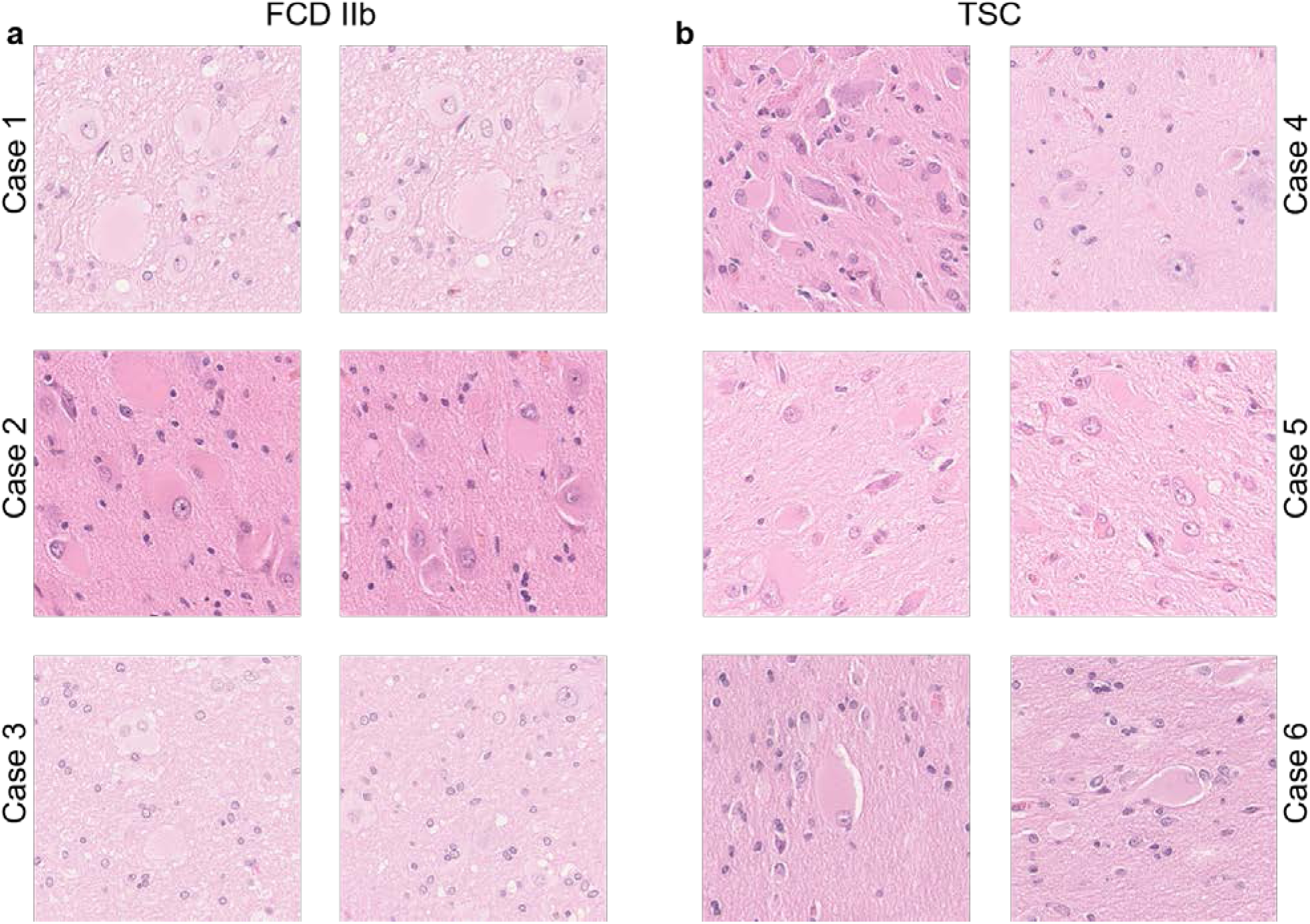
High power fields of FCD IIb and TSC. **a|** Three example cases with 2 high power fields (40x magnification) of FCD IIb. **b|** Three example cases with 2 high power fields of TSC in

In this study, we present a proof-of-concept deep learning approach to classify FCD IIb and TSC and to visualize the underlying distinguishing features which at the moment are not reliably discernible by pathologists in hematoxylin and eosin-stained (H&E) slides. This process is possible by computing guided gradient weighted class activation maps (Guided Grad CAMs) which represent the weighted pixel-level classification results which in turn mark the important histomorphological features the CNN uses to distinguish these entities. In addition, we implemented a custom slide review platform for whole slide image visualization and on demand classification.

Our approach might be a powerful concept for classifying and analyzing difficult to diagnose pathologic entities.

## Materials & Methods

### Dataset and Region of Interest

To train and evaluate our CNN, H&E tissue slides of 56 patients, who had undergone epilepsy surgery and were diagnosed at the European Neuropathology Reference Center for Epilepsy Surgery, were collected. The samples were subsequently digitized using a Hamamatsu S60 scanner. Overall the dataset consisted of 141 WSI from 56 patients, 28 patients with FCD IIb and 28 patients with genetically confirmed TSC. The whole dataset was divided into 50 cases used for training and validation along with 6 cases as an independent test-set to evaluate the model’s performance. We ensured that a patient was either in the training-and validation-set or the unseen test-set. The WSIs of our dataset were reviewed by two expert neuropathologists of the European reference center for epilepsy in Erlangen using the 2011 ILAE classification of focal cortical dysplasia [18]. The region of interest (ROI) on an individual slide was defined as areas with high balloon cell or giant cell counts along with the surrounding white matter and deep cortical layers. These ROIs were extracted at 20x magnification and cropped into smaller tiles of 2041×2041 pixels, using QuPath [19], to further preprocess and feed into our model.

### Convolutional Neural Network Architecture

A VGG16 CNN architecture pre-trained on ImageNet was implemented [3], using the open-source Python packages Keras [20] with TensorFlow backend [21]. VGG16 was chosen, as it yielded the best results with the least overfitting on a small training- and validation-subset out of a couple of state-of-the-art network architectures including NasNetMobile, Xception, DenseNet121 and ResNet50 [22–25]. The basic network architecture was not changed and consisted of 1 Input-Layer, 5 Convolutional Blocks each ending in a MaxPooling2D-Layer merging into a custom top layer beginning with a GlobalMaxPooling2D-Layer followed by 2 Blocks of Batch Normalization [26], Dropout (0.5, 0.5) and Dense-Layers merging into the fully-connected output layer (SoftMax-activation to produce individual output probabilities); (Figure 1.c).

### Preprocessing and Data Augmentation

Image preprocessing is an important step in every computer vision task to augment the number of samples, to prevent overfitting, and to support the model against invariant aspects that are not correlating with the label [27, 28]. In our approach, we mixed our novel random-rotate-zoom technique with classical image augmentation techniques. The initial 2041×2041 ROIs were cropped in 20x magnification. From these ROIs new sub-samples with random zoom and rotation were generated, resulting in 0.1 to 2x scaled sub-samples of the initial tile as shown in (Figure 2.a: Tile extraction and random-rotate-zoom Example). These sub-samples were then normalized and resized to obtain 300×300 pixel images. The 300×300 images were additionally augmented using the open-source, python-based library, imgaug [29] with a random composition of shear, blur, sharpen, emboss, edge detect, dropout, elastic transformations and color distortion including contrast adjustments, brightness changes, permutation of hue (all augmentations applying to either the whole image or an area of the image) (Figure 1.b). This process is implemented through a custom keras image generator. This image generator streams 50 training images, randomly generated, of every tile as an input into the CNN, using the described preprocessing method. By means of such procedures, there was no need to save any additional images to disk and, with random permutations on every training epoch, we maximized the learning efficacy and robustness of the neural network (Figure 4.b).

### Training and Evaluation

Training was performed with a batch size of 128, using the Adam-Optimizer [30] and a cyclic learning rate (cLR) [31] oscillating between 10^−8^ and 10^−3^ every quarter epoch, with a schedule to drop the cLR if validation-loss did not improve for 10 epochs. Training performance was controlled using accuracy, loss and area under the curve (AUC) as metrics where plotted every epoch. As a first step, base layer weights were frozen, only training the custom top layer with a cLR (10^−3^ – 10^−5^). In a second step, the whole model was trained including the base layers with a very low cLR (10^−6^ – 10^−8^), thus maintaining the basic image-classification patterns of the pre-trained model and prevent overfitting. Model parameters were saved every reduction of validation-accuracy and the best parameters were used for predictions on the unseen test-set.

We further evaluated model performance with 10fold cross-validation and a training- and validation-split of 0.9, while maintaining the original case-distribution and without having any training- and validation-slide overlap.

To evaluate model performance on the unseen test-set, images were generated using our random-rotate-zoom technique with 100 iterations on every test-tile, which were then individually predicted and averaged to obtain the prediction of a WSI. In the next step, the predictions on multiple WSIs of one case were averaged to obtain the prognosis for the whole case (Figure 2.c: Prediction Process). To further assess testing-performance the classification results were evaluated by accuracy, adjusted geometric mean (AGM), area under the receiver operating characteristic curve (AUC ROC), sensitivity, precision and F1 score (harmonic mean of precision and sensitivity).

### Visualization

Gradient-weighted class activation maps (Grad-CAM) and Guided Grad-CAM have shown to be useful tools to understand how the model is analyzing the images and revealing the features of relevance for the classification-task [32, 33]. Grad-CAMs are a form of localization maps based on a given image passing through the trained model to generate the class-gradient, setting all other class-gradients except the class of interest to zero and backpropagating the rectified convolutional features to compute a Grad-CAM [33]. In dependence of the convolutional layer different degrees of detailed spatial information and higher-level semantics are displayed. According to the original paper we expected the penultimate convolutional layer (block5_conv3) to have the best visualization results for our purpose. This Grad-CAM can be visualized using different colormaps to obtain a better understanding at which regions of a given image the model is focusing. Guided Grad-CAM is a combination of Grad-CAM’s “heatmaps” and guided backpropagation merged with point-wise multiplication to achieve pixel-level resolution of discriminative features [33]. We performed our studies based on Gildenblats’ open-source implementations of Grad-CAM and Guided Grad CAM for Keras [34], using our own models and adaptations of the code for our task-specific problems. This combination formed an optimal foundation to gain insight into which histomorphological features on region- and pixel-level, not discernible to the human eye, were relevant to distinguish between these two entities.

### Web application

We developed a custom web application for online slide review using Django and VueJs frameworks and openslide for WSI visualization. H&E specimens representing can be microscopically reviewed without any further information. Inside the application snapshots in 20x magnification can be taken on which regions a potential reviewer deemed important and submit the images from these regions via the web application. We then implemented an online classifier into the application which incorporated the trained model at the server-side. The online classifier worked as follows: An image could be taken directly from a WSI at 20x magnification and a predefined size at review and was subsequently stored on the server and classified by the model at the server-side. This way we assured the image fulfills certain quality criteria prior to being predicted by the model to obtain an averaged diagnosis over all the ROIs of one potential review-participant (Figure 3).

**Figure 3.**
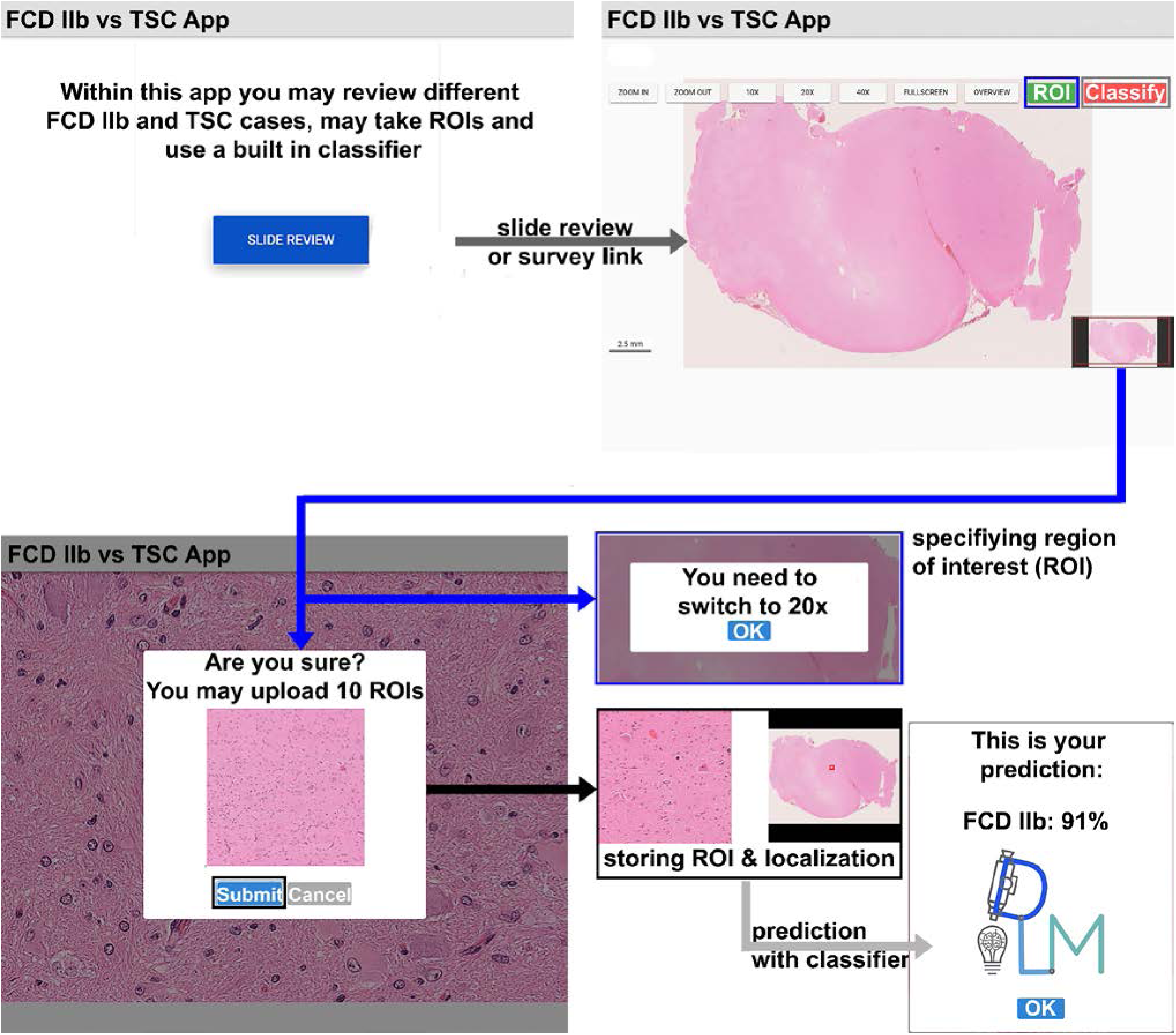
Slide overview of our self-developed slide review and prediction platform. Home screen with short introduction following the black arrow to enter the slide review. ROI selection and quality control (following the blue arrow), you can only screenshot ROIs in 20x magnification. Storing ROI and ROI localization on a server (2^nd^ black arrow). Prediction with random-rotate-zoom on all stored ROIs of one reviewer, with averaged prediction (grey arrow).

### Hardware

We implemented our approach on a local server running Ubuntu (18.04 LTS) with one NVIDIA GeForce GTX 1080Ti and one NVIDIA Titan XP, AMD CPU (AMD Ryzen Threadripper 1950X 16x 3.40GHz), 128Gb RAM, CUDA 10.0 and cuDNN 7.

## Results

### Validation and Test Performance

All CNNs were trained to classify the ROIs containing balloon cells and giant cells of FCD IIb and TSC-tuber, respectively, as well as surrounding tissue. First, we performed a study to determine which model to use for our classification task ranging from VGG16 to models with more trainable parameters, i.e. ResNet50, DenseNet121, NasNetMobile and Xception. We evaluated all those models on a small training- and validation subset of our whole dataset, with validation accuracy ranging from 75% (Xception) to 92% (VGG16) after 40 epochs (Supplement 1.b). We decided to implement VGG16 for our approach as it yielded the best validation results with little overfitting on the validation-subset, low training times and good architecture to visualize. In the next study, we compared our random-rotate-zoom preprocessing method with a direct ROI extraction of 300×300 pixel tiles on another training- and validation subset to determine if our approach improves classification performance and prevents overfitting. The results showed the superiority of the random-rotate-zoom technique over direct ROI extraction as validation accuracy was higher (Supplement 1.a).

Based on these studies we built our final VGG16 model with a custom top layer for extended Batch Normalization with random-rotate-zoom preprocessing and additional data augmentation (Figure 1.c). To assess the whole training set and select the best performing model for our final prediction, we evaluated our models via 10-fold cross-validation. The overall cross validation-performance is shown in Figure 4.a with a validation accuracy averaging at 94% (91%-97%) and an AUCROC averaging at 0.91 (0.89-0.95). During training validation accuracies mostly stayed above training accuracies and validation loss stayed below training loss values, indicating little to no overfitting on the training dataset.

**Figure 4.**
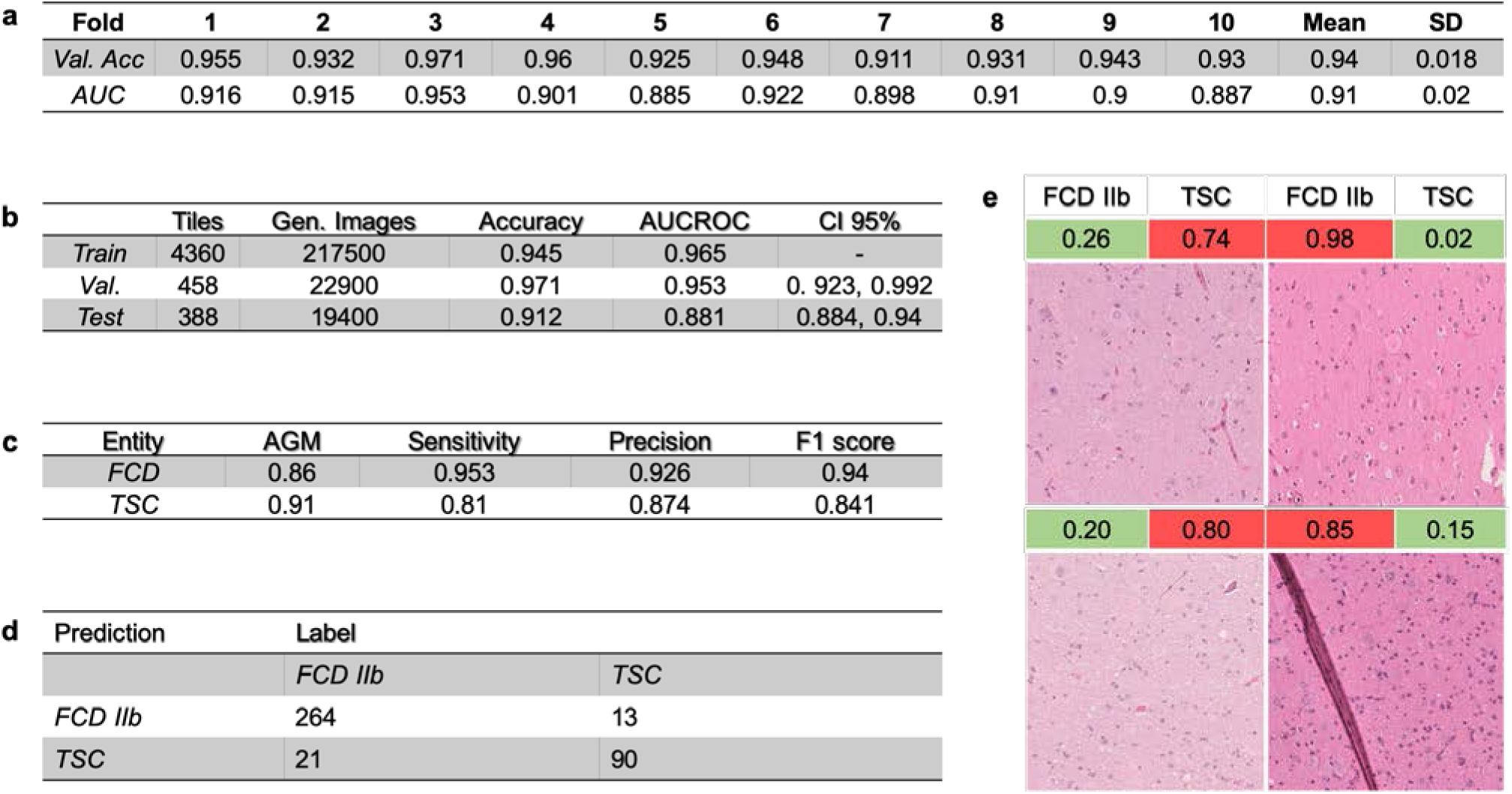
Data distribution and performance overview. **a |** Average predictions for each cross-validation fold with overall mean and standard deviation (SD). **b** | Data distribution as well as comparing view of training, validation and testing performance of our best performing model. **c** | Performance metrics for our best performing model in 10-fold crossvalidation on the unseen test set (average geometric mean (AGM)) **d** | Confusion matrix for the best performing model on the unseen test set on tile level. **e** | Most confident wrongly classified tiles for FCD IIb and TSC.

The best performing model in cross-validation was picked to classify the unseen test set. Scoring an overall accuracy of 91.2% on the tile level while not misclassifying a single case in our unseen test set (Figure 4.c). Additional performance metrics while testing are shown in Figure 4.c. The confusion matrix for the single tile predictions on the hold-out test set is shown in Figure 4.d. The results indicated a good overall performance for a classification task not easy to accomplish even for expert neuropathologists. To analyze problems and pitfalls of the trained CNN resident neuropathologists reviewed the most confident wrongly classified tiles (tiles with a high accuracy for the wrong label) to depict possible disruptive factors. The inspection of these tiles showed that some present folding artifacts (2/34) and stripy artifacts (6/34) due to tissue processing as well as some areas being slightly out of focus. Thereby making it more difficult to classify on a per tile basis (Figure 4.e).

### Model Visualization

We further investigated morphological features relevant to the classification task of both FCD IIb and TSC using Grad-CAM and Guided Grad-CAM. A total of 10000 Grad-CAMs and Guided Grad-CAMs heatmaps were generated and reviewed by three resident neuropathologists. Although ROI’s extracted via random-rotate-zoom reveal better classification results for visualization purposes a stable magnification level turned out to be better for the generation of accurate heatmaps (data not shown). This result is expected as the model can fit more precise to a single magnification level but generalizes worse when tested on the unseen test set.

The analysis of the generated Guided Grad CAMs revealed matrix reaction as an important feature distinguishing between cortical tuber and FCD IIb. In TSC patients the matrix reaction was fibrillar and strand-like throughout our visualized test-set of 10000 heatmaps (Figure 5.a). In contrast, the matrix reaction in FCD IIb specimens was diffuse and granular. Another new feature was that astrocytes and their nuclei played an important role in the morphological distinction between FCD IIb and TSC. Smaller nuclei of astrocytes with more condensed chromatin were a hallmark of FCD IIb (Figure 5.b), while larger nuclei of astrocytes with uncondensed chromatin structure were mainly found in TSC (Figure 5.b).

**Figure 5.**
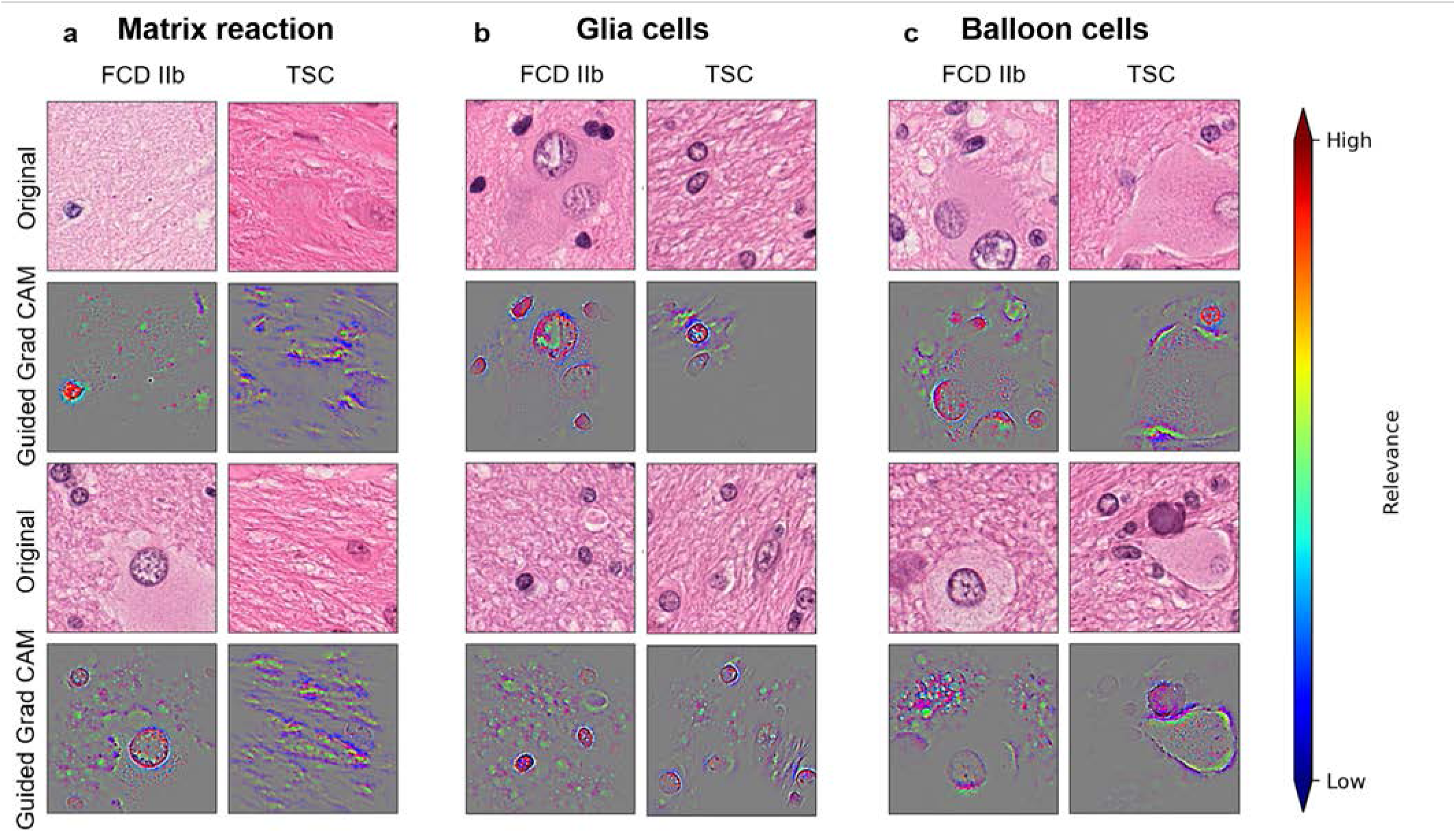
Visualization of representative Guided Grad CAM results matched with the original image tiles. **a** Matrix reaction, **b** Glia cells and **c** Balloon cells as seen by our best predicting model. Relevance scale is plotted with red being most confident and blue the least confident. Grey areas are unimportant to the prediction of the given class.

Surprisingly balloon cells were itself hardly used for the distinction of these two entities. The CNN often focused on the cytoplasm, cell wall and some chromatin sprinkles of balloon cells in TSC patients. An interesting finding in TSC patients were halo artifacts around balloon cells occurring in the majority of our dataset (Figure 5.c).

It is important to note that the CNN, even in images with artifacts like marker or small empty areas, did not target these structures to classify the tile.

## Discussion

We developed a deep learning approach to help diagnosing difficult-to-classify histopathologic entities and used CNN to extract novel distinguishing features. We then developed a new web-based application for histopathology diagnostics with a built in classifier.

At first, we evaluated different state-of-the-art model architectures to identify the most suitable for our purpose in terms of a) best classification results, b) least overfitting and c) best to visualize. Amongst all evaluated network we have chosen VGG as it accomplished these needs (see results). It is interesting to note, however, that models with higher network parameter counts and more complex architectures overfitted the given training data. In addition, VGG16 visualization through Guided Grad-CAMs has recently been used as a successful application in medical research, because it makes visualization at the pixel level possible [35–37]. The next step was to augment our use-case data set through appropriate preprocessing. Small datasets are of major concern for deep learning tasks and likely result in overfitting to the given training data and yielding inaccurate results of the independent test set [38]. We could show the benefit of effectively multiplying our data by a new random-rotate-zoom technique in addition to classical image preprocessing from direct patch extraction for the given task. This protocol was influenced by daily histopathology practice using different optical zoom ranges of the microscope to extract all available histomorphological information. Further and independent work is needed, however, to confirm the benefit of such preprocessing pipeline over direct patch extraction for deep learning tasks in digital pathology. Another important goal was to extract classifying features from our model using the Guided Grad-CAM approach. Patterns of matrix reaction and a halo artifact around balloon cells were novel and not yet described. The feature of condensed nuclei of astrocytes in FCD IIb compared to the uncondensed and bigger nuclei of astrocytes in TSC confirmed prior studies of the role of astrocytes in TSC and represented also a stable histomorphologic correlate in H&E stained sections [39, 40].

### Different limitations and possible solutions moving into the future

A well-recognized obstacle in digital pathology represents batch effects including variation in staining intensity or fixation artifacts [10, 41]. We contained such batch effects in our input data through hand-picked ROIs and normalization. However, more sophisticated H&E normalization standards needed to be developed to allow a comprehensive application of deep learning for the large spectrum of disease conditions [42, 43]. In particular, integration of cell-type-specific brain somatic gene information into disease classification will advance inter- and intra-observer liability of histopathology diagnosis as well as better understanding of underlying pathomechanisms [44].

Our dataset was small with respect to deep learning standards, especially when comparing datasets collected by Imagenet or OpenImage with a compilation of million images and thousands of unique samples per class when training deep learning architectures [45, 46]. But this will be an impossible task when studying rare brain diseases. Our dataset of 56 cases is extraordinarily large and was collected with the help of the archives of the European Epilepsy Brain Bank [47]. Such sample numbers exceed what most pathologists will see in their lifetime. Hence, we call for multi-center collaborations to obtain big enough datasets and to develop open-access online tools for consultation if a certain disease is suspected. Furthermore, our approach passed many previous endeavors in size of the dataset and sophistication of preprocessing. Our approach may support the diagnosis of rare MCD entities in regions of the world with no genetic testing available [48, 49]. Despite the fact that such online tools are not yet approved for diagnostic use, we think it will help to disseminate web-based digital pathology tools into regions of the world where genetic testing or advanced neuropathology expertise is not a common or even available standard.

In conclusion, our study demonstrated the successful use of deep learning in the diagnosis of histomorphologically difficult-to-classify MCD entities. Morphological features learned by the system and relevant for the classification of FCD IIb and cortical tuber were then visualized. These results are promising and will help to amplify CNN visualization and deep learning methodologies in the arena of digital pathology.

## Availability and implementation

Project Home Page: https://github.com/FAU-DLM/FCDIIb_TSC

## Acknowledgments

The present work was performed in fulfillment of the requirements of the Friedrich-Alexander Universität Erlangen-Nürnberg (FAU) for obtaining the degree ‘Dr. med.’ of Joshua Kubach. We would like to thank NVIDIA for the donation of a Titan XP.

## Supplement

**Supplement 1.**
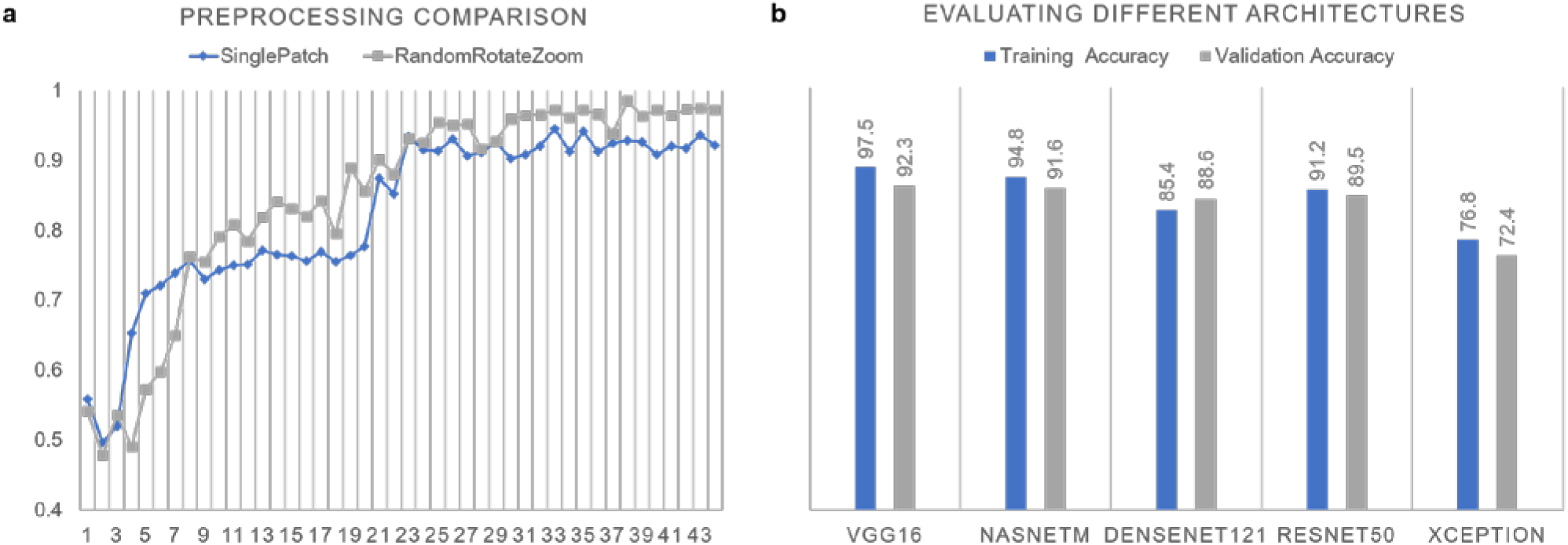
Selection of the right Preprocessing and Model-Architecture. **a**| Comparison of Single Patch Extraction (300×300 Tiles) versus random-rotate-zoom (2000×2000 Tiles with subsequent random sub-tile extraction & resizing to 300×300). **b**| Comparison of VGG16, NasNetMobile, DenseNet121, ResNet50 and Xception on a subset of our whole dataset with 40 training epochs for each model.

